# Changes in capture availability due to infection can lead to correctable biases in population-level infectious disease parameters

**DOI:** 10.1101/2022.09.23.509235

**Authors:** Iris A. Holmes, Andrew M. Durso, Christopher R. Myers, Tory A. Hendry

**Affiliations:** Cornell Institute of Host Microbe Interactions and Disease, Cornell University, Ithaca, NY, USA; Department of Microbiology, Cornell University, Ithaca, NY, USA; Department of Biological Sciences, Florida Gulf Coast University, Fort Myers, FL, USA; Center for Advanced Computing & Laboratory of Atomic and Solid State Physics, Cornell University, Ithaca, New York, USA

## Abstract

Correctly identifying the strength of selection parasites impose on hosts is key to predicting epidemiological and evolutionary outcomes. However, behavioral changes due to infection can alter the capture probability of infected hosts and thereby make selection difficult to estimate by standard sampling techniques. Mark-recapture approaches, which allow researchers to determine if some groups in a population are less likely to be captured than others, can mitigate this concern. We use an individual-based simulation platform to test whether changes in capture rate due to infection can alter estimates of three key outcomes: 1) reduction in offspring numbers of infected parents, 2) the relative risk of infection for susceptible genotypes compared to resistant genotypes, and 3) change in allele frequencies between generations. We find that calculating capture probabilities using mark-recapture statistics can correctly identify biased relative risk calculations. For detecting fitness impact, the bounded nature of the distribution possible offspring numbers results in consistent underestimation of the impact of parasites on reproductive success. Researchers can mitigate many of the potential biases associated with behavioral changes due to infection by using mark-recapture techniques to calculate capture probabilities and by accounting for the shapes of the distributions they are attempting to measure.

## Introduction

Emerging infectious diseases, driven by climate change, introduced species, and other anthropogenic disturbances, are a conservation concern for many animal populations (Lafferty and Gerber 2002; Smith et al. 2009; Morand 2020). Detecting pathogen-induced selection in natural populations is key to managing the threat they pose. Pathogens can impose strong fitness consequences on hosts, potentially reducing population growth and long-term stability (Maslo and Fefferman 2015; Iverson et al. 2016; Campbell et al. 2018). In addition, selection by parasites on hosts can lead to conservation-relevant evolutionary changes if resistance phenotypes are heritable (Prugnolle et al. 2005; Fumagalli et al. 2009; Kariuki and Williams 2020). For example, a population that can quickly adapt to a novel disease may require less active management than a population that cannot (Spielman et al. 2004; Mikheyev et al. 2015). Conversely, strong selection toward infection-resistant genotypes may lead to reduced population-level genetic diversity (Jordan et al. 1998; Lenz et al. 2016). However, estimating the strength of selection or its correlates in wild populations is logistically challenging (Chao 1989; McDonald and Amstrup 2001; Grimm et al. 2014). Here, we use a simulation approach to identify scenarios in which estimates of parasite-induced selection may lead to spurious conclusions. We also demonstrate strategies that can be used to mitigate the risk of incorrectly identifying correlates of selection when none is occurring or incorrectly ruling out selection when it is occurring.

A central challenge in connecting parasite infection to host fitness in natural populations is that infection can alter capture rates of hosts (Benton and Pritchard 1990; McPherson et al. 2012; Garamszegi et al. 2015). Mark-recapture approaches are the state of the art in accounting for differences in capture likelihood between subgroups within a focal population. Robust design mark-recapture methods use the capture history of individual animals over several bouts of sampling. Capture histories are a record of each time an individual is sampled or not during sampling bouts. If an individual is sampled in bouts one and three, it can be safely assumed it was present during bout two but was not captured. The ratio of successful captures to misses can then be used to calculate the capture rates for pre-identified subsections of the study population (Pollock et al. 1990; Nichols 1992; Willson et al. 2011).

Parasites impact capture availability in a variety of ways. Parasites can reduce escape performance (Benton and Pritchard 1990; Schall 1990), causing a reduction of host activity and thereby a reduction of capture rates of infected individuals (Bass and Weis 1999; McPherson et al. 2012; Koprivnikar and Penalva 2015). Conversely, the energetic demands of parasite infection could drive the host to greater foraging efforts, thereby increasing availability for capture (Benton and Pritchard 1990). Some parasites manipulate host behavior to increase risk of predation, allowing the parasite to move from an intermediate to a definitive host (Levri and Lively 1996; Lagrue et al. 2007), which could increase capture rates of infected hosts. In addition to differences in capture probability, simple sampling error can impact estimates of key outcomes. This is particularly true when sampling from bounded distributions, when a parameter can vary freely in a specific range of values but not outside those values. These distributions are common in biology, when parameters are often constrained to positive numbers, for example the concentration of a protein (de Franciscis et al. 2014), the population size of an animal (Cai and Geritz 2020), or the frequency of an allele in a population (Kimura 1957).

There are several correlates of pathogen-driven selection that can be measured in wild host populations. When a biologically plausible resistance allele has been identified, quantifying changes in genotype frequencies between generations within a population can provide strong evidence of selection occurring (Westerdahl et al. 2004; Thrall et al. 2012). In addition, determining whether differing rates of infection are associated with different alleles or genotypes can provide evidence of selection (Langefors et al. 2001; Froeschke and Sommer 2005; Dionne et al. 2009; Sin et al. 2014). Demonstrating differential reproductive success based on infection state is also critical to showing that selection may be acting, as some parasites do not impact lifetime reproductive success and so cannot drive selection (Schall 1983; Gustaffson 1994; Zylberberg et al. 2015).

One frequently studied family of vertebrate genes that can confer parasite resistance are the Major Histocompatibility Complex, or MHC. MHC proteins are responsible for recognizing pathogens and starting the adaptive immune response cascade (Kaufman 2018). High MHC diversity can increase fitness by allowing an animal to mount immune responses to a broader variety of pathogens (Agudo et al. 2012; Radwan et al. 2012). However, in some systems a specific MHC allele will confer the strongest fitness benefit (Froeschke and Sommer 2005; Wroblewski et al. 2015). Other gene families that are less studied than MHC, but may experience similar switches between directional and balancing selection due to pathogen pressure, include the immunoglobulin A genes as well as scent and taste receptors, which play a role in recognizing pathogens in many tissues in the body (Sumiyama et al. 2002; Shi et al. 2003; Seixas et al. 2012; Carey and Lee 2019; Harmon et al. 2021).

Here, we establish a simulated population based loosely on the ecology of lizard-malaria systems. In such systems, parasite infection reduces host lifetime reproductive success but does not shorten host lifespan (Dunlap and Schall 1995; Eisen 2001), a common pattern for sublethal parasites (Dyrcz et al. 2005; Marzal et al. 2005; Hillegass et al. 2010). We examine both heterozygote-advantage and resistance-allele advantage scenarios. We quantify the impact of random subsampling and biased detection on our ability to estimate three outcomes of interest: 1) the fitness impact of infection, 2) the relative risk of infection of different host genotypes, and 3) changes in allele frequency in the population over a single generation.

## Methods

### Simulation framework

We simulated a diploid host population with two alleles at a single locus in the R v4.4.1 scripting environment (R Core Team 2021). Each simulation had at least one genotype that conveyed resistance to a pathogen. In the ‘heterozygote’ runs, heterozygotes had low infection risk, while both homozygous genotypes had higher infection risk. In the ‘resistance allele’ runs, carriers of a resistance allele, whether heterozygous or homozygous, had lower infection risk. Individuals became infected or not in a Bernoulli trial, with the probability of infection determined by their genotype and a potential additional stochastic component drawn from a uniform distribution between zero and one.

After infection, parents were randomly sampled from the population. The number of offspring assigned to each pair was sampled from a Poisson distribution with a set mean. For each infected parent, the mean of the Poisson distribution was altered down by a set proportion of the mean expected number of offspring from two uninfected parents. The parents’ genotypes were randomly subsampled to create gametes for offspring genotypes. Once offspring had been generated for a set number of breeding pairs, the offspring pool was subjected to infection as described above and added to the full population.

We conducted five sampling events in which we randomly drew a set number of individuals from the full population. We apply a Cormack-Joly-Seber model implemented in the R package ‘marked’ to these data to identify whether differences in capture rates between infected and uninfected individuals could be detected (Laake et al. 2013). We performed three separate runs for each set of parameter values: a control run in which all individuals had the same capture probability, an ‘increased rate’ run in which infected hosts had a higher capture rate, and a ‘reduced rate’ run in which infected hosts had a lower capture rate. We randomly selected an individual and used the applicable capture probability in a Bernoulli trail to determine whether that individual was captured or not. We continue the process until we reached our designated sample size. We then applied the CJS model to calculate capture probability for infected and uninfected individuals in these samples.

For each outcome of interest, we performed 200 runs of the model for each of three capture rate scenarios. We randomly drew 200 values from two different uniform distributions for a parameter that described the proportional difference in capture rates between infected and uninfected hosts. One distribution was between 0.1 and 0.9 (reduced rate), and one between 1.1 and 1.9 (increased rate), while our control runs had no difference in capture rate between infected and uninfected hosts. For each question, we perturbed a second simulation parameter to test the impacts of differing capture rates across a range of possible scenarios. For our offspring penalty simulations, we perturbed the parameter that controlled the expected proportion of offspring lost due to infection in the parents. We selected values from a uniform distribution between zero and one for this parameter. For our relative infection risk and allele frequency change simulations, we drew values from a uniform distribution between zero and one for the parameter controlling the degree to which the genotype of the host predicts infection risk. We recorded the results of these simulations and uploaded the results, along with the code, on Zenodo (DOI: 10.5281/zenodo.6639310). The values of the statistical tests in the results section are derived from these 200 recorded runs. For the sake of visual clarity, the figures are based on the first 50 runs in the outputs.

### Statistical tests

For each of our three outcomes of interest, we applied the same set of statistical tests to the values derived from the full population compared to the control samples, the increased rate samples, and the decreased rate samples. For the genotype relative risk and allele frequency change simulations, we performed separate analyses on the heterozygote and resistance allele runs. We use a slope test implemented in the R package ‘smatr’ to identify whether the regression of the increased rate, reduced rate, or control samples measured against the true values had a slope significantly different than one to one (Warton et al. 2012). To test for greater spread of points in the different-rate samples relative to the true values and control samples, we use a Fligner test implemented in the R package ‘stats’ (R Core Team 2021).

Finally, we used two approaches to determine whether the Cormack-Joly-Seber mark-recapture model correctly identified runs with greater sampling bias imposed by the differences in capture rates. We performed t-tests in base R between the control and both infection-dependent capture rate outcomes using the difference between the capture rate value for infected and uninfected individuals. This test showed whether the CJS capture rate values correctly identified altered capture availability in our simulation runs. Second, we found the residuals of a linear regression between the values for the parameter of interest from each captured sample and the full-population values. We regressed the residuals against the absolute difference in the CJS model capture probability values for infected versus uninfected hosts. If the CJS model correctly identified the magnitude of the bias in our parameter values, we would expect a positive correlation between these values.

## Results

### Fitness impacts of infection

We used slope tests to detect whether differences in capture rates between infected and uninfected hosts influenced the accuracy of the estimation of fitness reductions in infected parents. When we regressed raw values of the difference in offspring number from the control sample relative to the full population, the sampled values dramatically underestimated the magnitude of the difference between reproductive success of the infected and uninfected parents (Figure 2A, slope=0.15, p=0). When we corrected all values by dividing the difference by the mean number of offspring for uninfected parents (Figure 2B), the value of the slope of the regression of the control samples against the full population values was much closer to one (slope = 1.16, p=9.24 × 10^−6^). The underestimation was likely due to the asymmetrical sampling space available. Parents with no offspring will always have their success ‘correctly’ detected by sampling, while the success of parents that do reproduce will be underestimated when some offspring are not captured.

**Figure 1:**
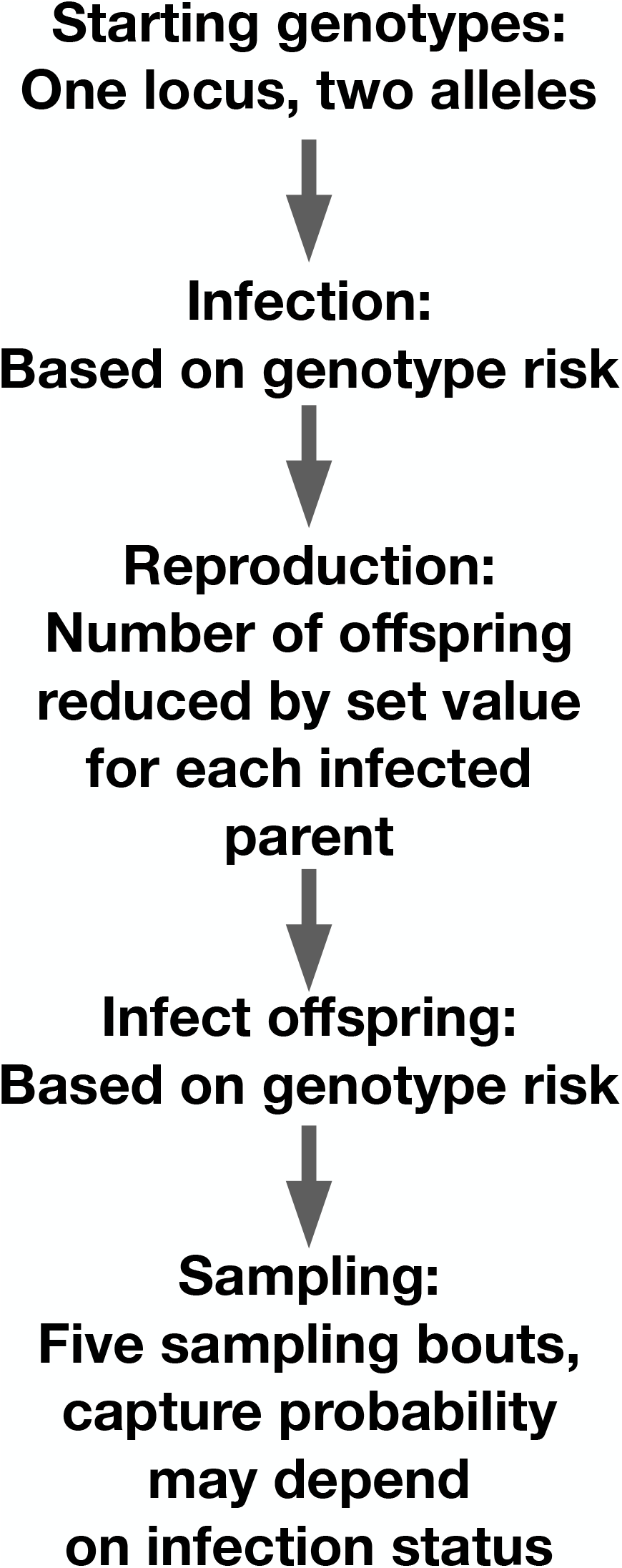
Schematic of our simulation setup.

**Figure 2:**
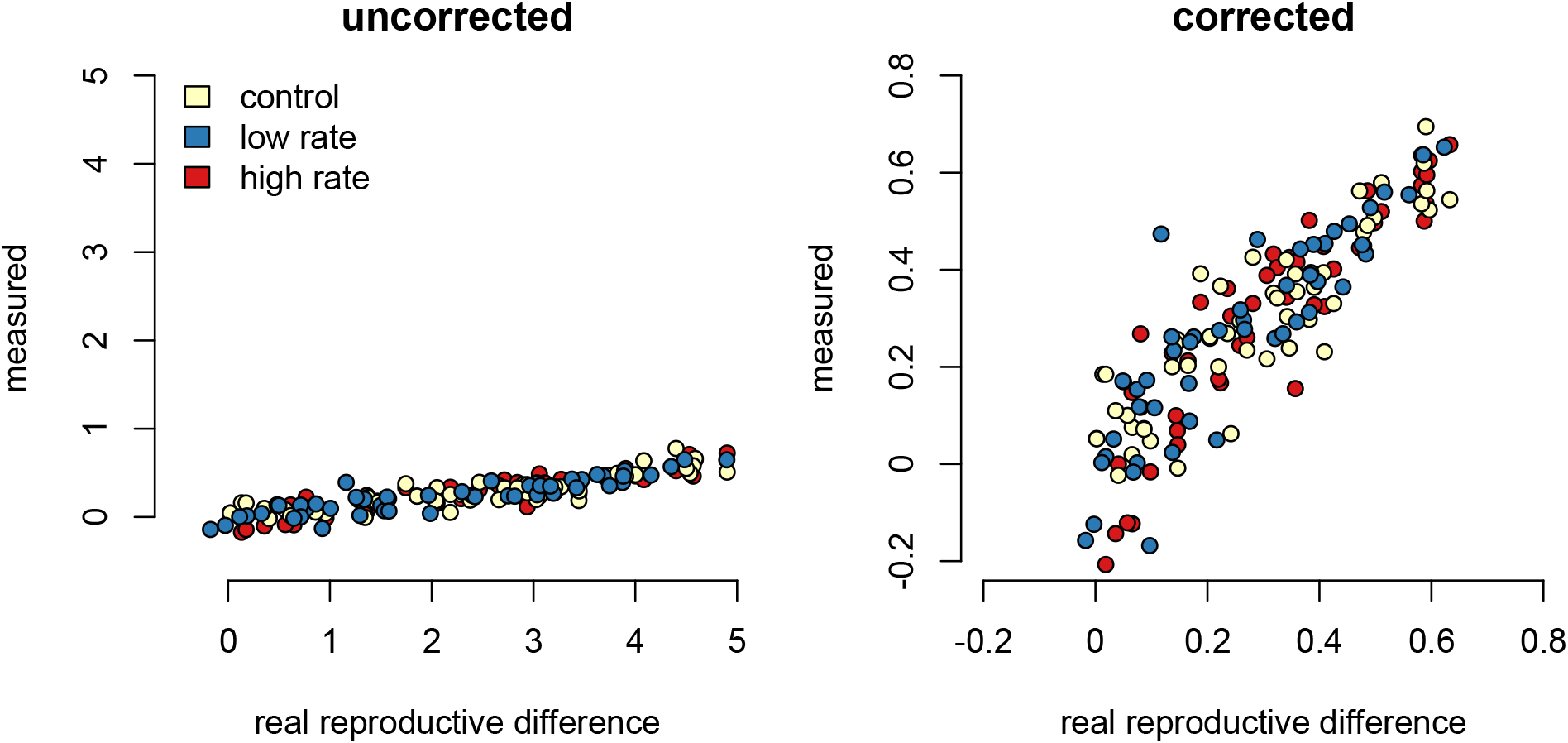
Sampling effects on detecting the impact of infection on reproductive success. Point colors represent different host reactions to parasite infection: 1) control samples in which infected and uninfected individuals have equal capture probabilities, 2) reduced rate samples in which parasitized individuals have a lower capture probability, and 3) increased rate samples in which parasitized individuals have a higher capture probability. Uncorrected sampling underestimates the impact of infection due to the uneven bounding of the parameter space. Standardizing the measured values by the mean number of offspring for uninfected parents reduces the sampling effects.

We found that slopes from corrected increased and decreased capture rate samples were not significantly different from one (increased p=0.960; decreased p=0.859). Since there is a stochastic component to infection for both parents and offspring, even susceptible parents with susceptible offspring are likely to have offspring in both infection categories. As a result, there are likely an adequate number of offspring in all infection categories to correctly detect relative reproductive success values.

To test whether sampling resulted in a larger variance of outcome values relative to the corrected true population values, we performed Fligner tests comparing the corrected reduced rate, increased rate, and control simulations to the true population values. Only the test of the control was significant (p=0.050), while the two differing rate samples did not have significantly higher variance than the real samples (increased p=0.103l; decreased p=0.094). This is likely due to the altered sampling rates clustering outcomes together, thereby counteracting the variability caused by random sampling effects.

T-tests successfully separated the differences in the capture rates of infected and uninfected individuals between control and varied capture rate runs (decreased rate p < 2.2 × 10^−16^; increased rate p < 2.2 × 10^−16^). To test the success of the CJS model on identifying specific runs with higher capture bias values, we first found the residuals of a linear regression of reproductive success differences from the captured versus the full populations. These measured the degree to which the outcomes of the simulation differed from the expected outcomes at a specific parameter value. Then, we found the absolute differences in the CJS capture probability values for the infected versus uninfected groups in each simulation run. We found the slope and R^2^ goodness of fit between the residuals and the capture probability differences. None of the three parameter values showed a significantly positive slope (increased p= 0.8189; decreased p=0.051; control p = 0.283), and all had small R^2^ values (increased R^2^ = -0.005; decreased R^2^ = - 0.014; control R^2^ = -0.0007).

### Relative infection risks of host genotypes

Regressing the relative risk measures from the heterozygote runs against the full population resulted in a slope close to one (slope = 1.04, p=0.002). The increased rate samples had a slope of 0.835, significantly lower than the control samples (p=0). The reduced rate samples had a slope of 1.56, significantly higher than the control slope (p=0) (Figure 3A). The resistance allele runs had a similar pattern, with the slope of the control sample regression being 1.00 (p=0.672) (Figure 3B). The increased rate sampling significantly underestimated the size of the relative risk between different genotypes (slope = 0.835, p=0), while the reduced rate samples significantly overestimated it (slope = 1.560, p=0). Results were similar for the resistance allele simulations (control: slope = 1.05, p= 6.43 × 10^−9^; increased: slope=0.755; p=0, decreased: 2.04, p=0).

**Figure 3:**
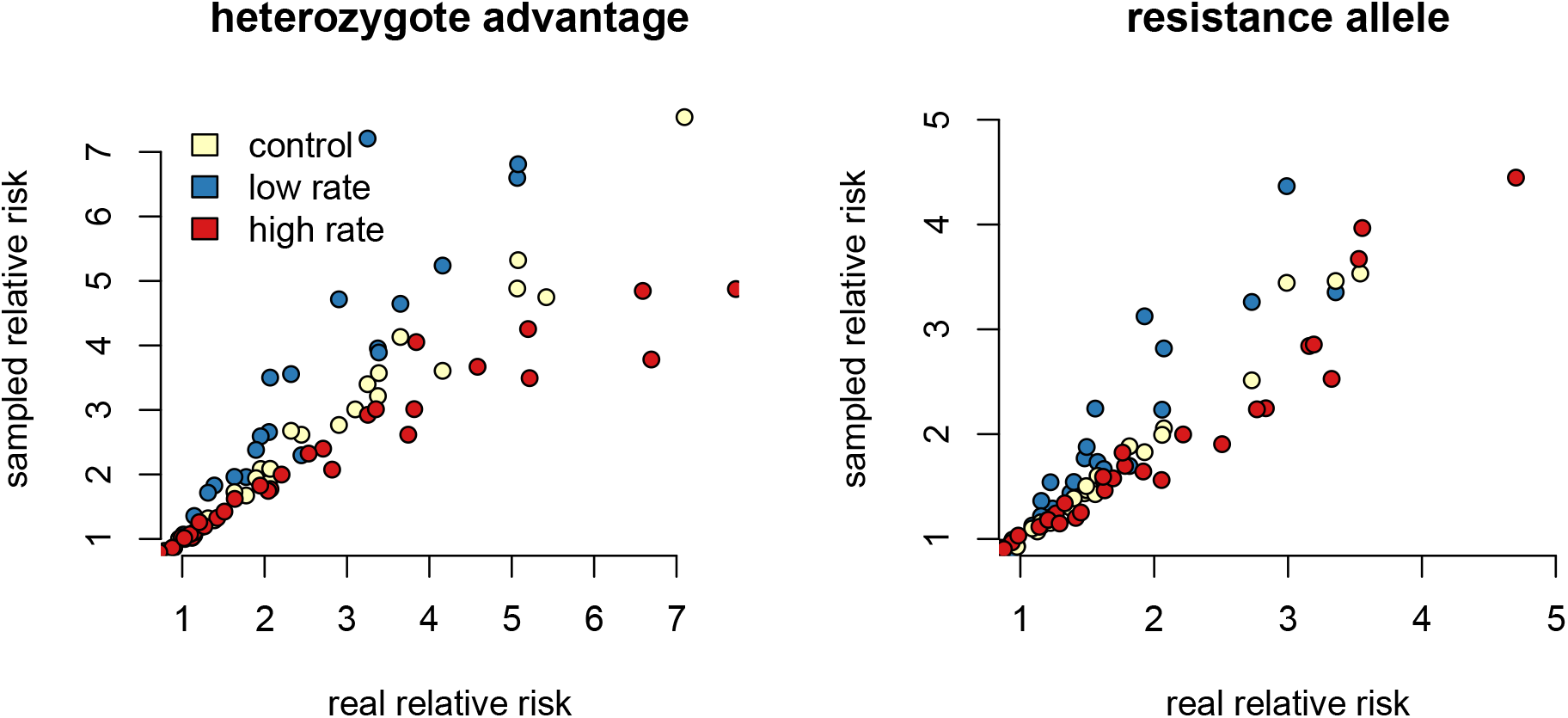
Differing capture rates impact estimates of relative infection risks of susceptible and resistant genotypes, in both the heterozygote advantage, and resistance allele scenarios.

In the decreased rate simulations, infected individuals with the resistant genotype will be the rarest sampled category. The relative risk calculation divides the proportion of infected individuals with the high-risk genotype by the proportion in the low-risk genotype. Failing to accurately measure the number of infected individuals in the low-risk genotype will inflate the denominator of the relative risk calculation, driving up the final value. The rarest sampled category is the most vulnerable to random sampling error. This produces the triangle-shaped distribution visible in Figure 3, in which some points in each increased and decreased rate samples fall near the control samples, but others deviate. Similar logic applies to the simulations in which infected individuals are more likely to be captured.

According to the Fligner test, the variability in points from the differing rate samples in the heterozygosity runs was significantly greater than the variability of the full dataset (increased rate p = 0.037; reduced rate p = 0.006). The variance in the control samples were not significantly different from variance it the full dataset (p=0.850). For the resistance allele runs, the pattern was similar, with differing rate samples being significantly different from the full dataset (increased p = 0.015; reduced p =0.005), while the control sampling was not (p = 0.889).

Linear models of residuals compared to capture probability values were significant for some parameter values. For the heterozygote simulations, the increased and decreased rate relationships were significant (increased: p =0.004, R^2^ = 0.037; decreased: p = 2.6 × 10^−5^, R^2^ = 0.081), while the control runs were not (p = 0.084, R^2^ = 0.010). The resistance allele simulations followed a similar pattern, although the increased rate samples did not show a significant correlation (increased: p = 0.109, R^2^ = 0.008; decreased: p = 5.09 × 10^−16^, R^2^ = 0.280; control: p=0.311, R^2^ = 0.0002). T-tests separated the differences between the capture rates of infected and uninfected individuals between control and varied capture rate in both simulations (p < 2.2 × 10^−16^ for all comparisons).

### Allele frequency change detection

In the heterozygote-advantage simulations (Figure 4A), allele frequency change calculated from the control samples showed a positive correlation with the true values (slope = 3.86, R^2^ = 0.891, p = 0). The increased rate and decreased rate simulations behaved similarly (increased: p=0, slope = 3.785; R^2^ = 0.879; decreased: p=0, slope = 4.024, R^2^ = 0.895). In the resistance allele simulations (Figure 4B), slopes were closer to one-to-one (control = 1.291; increased rate = 1.309; reduced rate = 1.245). Correlations looser but still significant (control R^2^ = 0.373; increased R^2^ = 0.415l reduced R^2^ = 0.368), and p-values were all significant (control p = 6.672 × 10^−8^; increased p = 8.179 × 10^−8^; reduced p = 1.355 × 10^−5^). Differing capture rate samples had significantly more variance in allele frequency change than the full population in both heterozygote-advantage (increased p = 2.2 × 10^−16^; reduced p = 2.2 × 10^−16^) and risk-allele (increased p = 0.003; reduced p = 0.02) simulations. Control samples in both simulation shad significantly higher variance than the full population (p = 2.2 × 10^−16^ for both).

**Figure 4:**
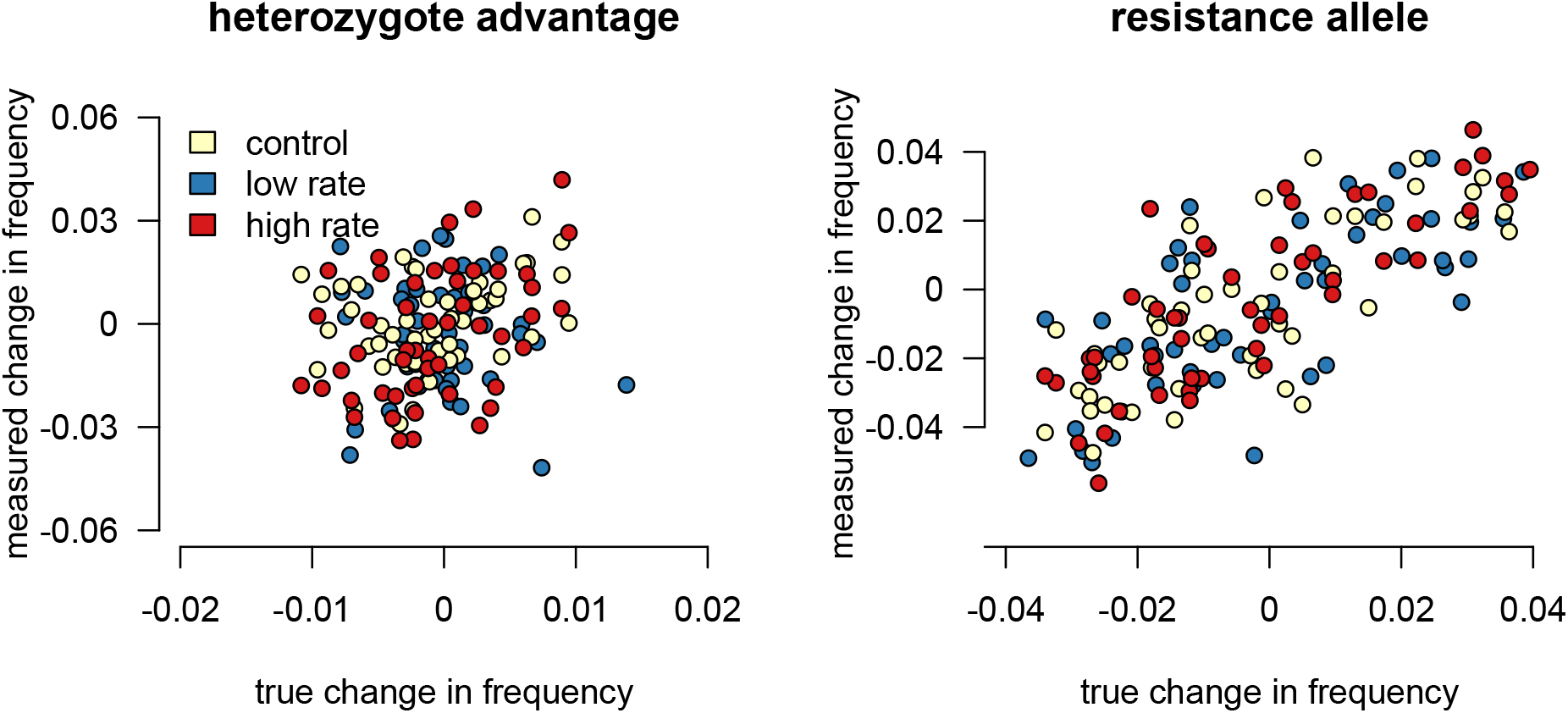
Sampling effects on measuring changes in allele frequency between generations. In the heterozygote-advantage case, no linear relationship exists between measured and true allele frequency changes, and measured changes can be double true changes. In the resistance allele simulation, a relationship does exist, but the sampled population can still overestimate the true allele frequency change.

For the heterozygote runs, none of the regressions of CJS capture rates against residuals reached the threshold of significance (increased: p=0.833, R^2^ = -0.005; decreased: p = 0.547, R^2^ = -0.003; control: p = 0.228, R^2^ = 0.002). The resistance allele simulations performed similarly (increased: p = 0.546, R^2^ = -0.003; decreased: p = 0.423, R^2^ = -0.002; control: p=0.832, R^2^ = - 0.004). As with all other comparisons, t-tests separated the differences between the capture rates of infected and uninfected individuals in all simulations (p < 2.2 × 10^−16^ for all comparisons).

## Discussion

Infection-induced capture bias and sampling effects impacted the reliability of our three ecoimmunological outcomes of interest. Differing capture rates can inflate estimates of the relative risk associated with susceptible genotypes (Figure 3), while sampling error can result in underestimating the impact of infection on reproductive success (Figure 2). Sampling error can also inflate measured changes in allele frequencies between generations, leading to overestimation of the impact of selection due to parasite infection (Figure 4). Mark-recapture statistics successfully identify runs in which infected and uninfected individuals have different capture rates. In one of our outcomes of interest, the relative risk of infection of different host genotypes, the difference in CJS capture rates were positively correlated with outlier values (Figure 5).

**Figure 5:**
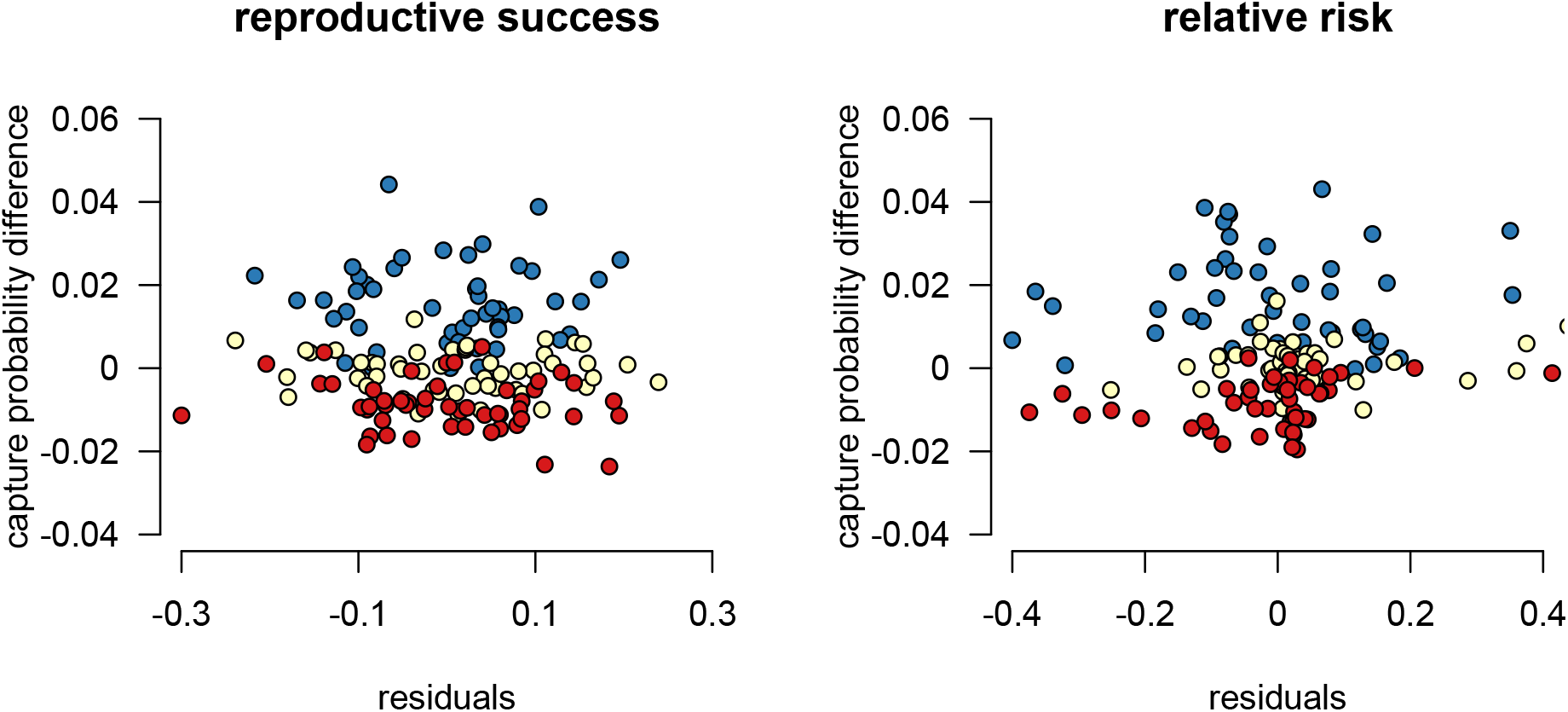
Robust design mark-recapture probability value differences successfully separate simulation runs in which infected and uninfected individuals have different underlying capture probabilities across our outcomes of interest. For measuring reproductive success differences, the residual values of the outcome of interest are not correlated with the success in detection of bias, while they are for detection of differences in genotype relative risk of infection.

Our simulations underestimated the impact of parasitism on reproductive success (Figure 2). We believe that this underestimate comes from the uneven bounding of the sampling distribution for this outcome (Kimura 1957; de Franciscis et al. 2014; Cai and Geritz 2020). An individual with no offspring will have their reproductive success “correctly detected” in every case because no offspring are available for sampling. However, highly successful parents are most likely to have estimates of their reproductive success far from their true value because they have more offspring in the population that can be missed. The bounded distribution can result in underestimation of the selective pressure of parasite infection if differences in reproductive success are left uncorrected. When an infection exerts selective pressure, uninfected parents should be the most reproductively fit. As a result, uninfected parents are the most likely to have their total success underestimated, which in turn dilutes estimates of selective pressure. Because the underestimates do not result from a difference in detection probability between infected and uninfected individuals, mark-recapture statistics are not helpful in identifying specific instances in which the issue is particularly acute. We find that adjusting for sampling effects by dividing the difference in reproductive success by the mean number of offspring captured from uninfected parents can nearly account for this source of error (Figure 2B). Future work may be able to generalize this approach or develop different normalization approaches.

Our measures of the relative risk of infection associated with susceptible genotypes were inflated when infected individuals were less likely to be captured and biased down when infected hosts were more available for capture (Figure 3). This outcome resulted from a combination of the altered sampling rates and sampling error. When the values of rare classes, such as infected individuals with resistant genotypes, drive a metric of interest, the metric is highly susceptible to random sampling effects within the rare outcome class. Because the alterations from the control group were driven in part by differences in capture availability, mark-recapture statistics were able to identify more-impacted simulation runs.

Sampling error more than differences in capture rates impacted our ability to correctly detect allele frequency changes, particularly in our heterozygote-advantage simulation (Figure 4). In our heterozygote-advantage simulations, true allele frequency changes in the population are less than half of the magnitude of the measured changes, and there is no significant relationship between the measured and the actual changes. In the resistance allele simulations, there is a relationship between measured and real allele frequency changes, and the amplitude of the allele frequency change can be more successfully detected from the samples.

Error in estimating parameters relevant to pathogen-driven selection on host populations can propagate into the estimation of a range of population-scale evolutionary scenarios. As with any organism, the pathogen population’s potential for adaptation is related to its size, which will be determined in part by the outcomes of host/pathogen coevolution. For example, the expected length of pathogen persistence in a host population is influenced by our focal parameters (Rand 1995; Fleming-Davies et al. 2015). Overestimating the relative risk of infection to susceptible hosts might lead to either the assumption that the host population will evolve toward resistance over short time frames (Lattorff et al. 2015; Vitale and Best 2019; White et al. 2021). Pathogen population size might then be assumed to be lower than reality (Kao 2006; Kerr et al. 2006), causing underestimation of the pathogen’s adaptive potential (Antolin 2008; Gordo et al. 2009; Ailloud et al. 2019). Such incorrect inferences could impact short and long-term conservation planning for disease management (Frick et al. 2017). Epidemiological conclusions, particularly for poorly understood emerging disease, can also be impacted by incorrect inferences about local selection dynamics (Keeling and Gilligan 2000; Ball et al. 2015; Britton and Scalia Tomba 2019).

In addition to evolutionary potential, within-population behavior is key to understanding a pathogen’s metapopulation dynamics. Several parameters of metapopulation models, such as propagule pressure and the expected longevity of a host or pathogen population, are impacted by our outcomes of interest. Like other organisms, pathogens can experience local extinction events. Pathogen persistence at a landscape scale is therefore dependent on their ability to migrate between subpopulations of their hosts (Soubeyrand et al. 2009). Even if the pathogen will eventually go locally extinct, a longer expected persistence will provide more opportunities to colonize naive host subpopulations and persist at the landscape scale (Laine 2004).

Metapopulation models are used to predict the spread and impact of emerging infectious diseases parameters (Keeling and Gilligan 2000; Ball et al. 2015; Britton and Scalia Tomba 2019), predict the evolutionary trajectory of virulence at the landscape scale (Thrall and Burdon 2003), and determine the likelihood that some host subpopulations will remain uninfected (Dijk et al. 2022).

Field studies have several sources of uncertainty that are not modeled here. First, false negative tests for pathogen presence can occur. Using mark-recapture techniques, the probability of non-detection of infection can be modeled jointly with imperfect detection of hosts (Jennelle et al. 2007; Conn and Cooch 2009; Cooch et al. 2012; Tersago et al. 2012). Second, imperfect detection of parent-offspring relationships can impact estimates of fitness. As with pathogen detection, uncertainty in genealogical reconstruction can be accounted for in empirical systems (Zhu et al. 2007; Lacy 2012; Wang 2017, 2019). While our simulations represent a best-case scenario, they point to broad patterns of bias that could be encountered in natural populations.

In summary, accounting for parameter estimation errors due to differences in capture rate and sampling error is a key concern in expanding landscape-scale host-pathogen evolution to a broad range of species. Mark-recapture techniques can be helpful in accounting for the ways that parasite infection can impact host behavior and therefore detection probability. Many mark-recapture studies are intended to detect survival probability of animals over relatively long time scales. These studies are time and resource intensive, because they require a large enough sample size of individuals to ensure that some can be recaptured throughout the study. However, we demonstrate in this paper that a single, week-long, bout of robust-design sampling has considerable value in accounting for detection bias between infected and uninfected individuals. Implementing this approach in the field could increase the accuracy of disease ecological sampling across many taxa of interest.

## Acknowledgements

This work was supported by the National Institutes of Health and National Institute of Allergy and Infectious Diseases Award T32Il45821.

## Notes

https://zenodo.org/record/6639310#.Yy4CTezMJJU

### Competing Interest Statement

The authors have declared no competing interest.

